# Assessing the Biosecurity Risk of Footwear as a Fomite for Transmission of Adventitious Infectious Agents to Mice

**DOI:** 10.1101/2024.11.03.621730

**Authors:** Michael B Palillo, Noah Mishkin, Mohamad Atmane, Jack A Palillo, Sebastian Carrasco, Cheryl Woods, Kenneth S. Henderson, Neil S Lipman, Rodolfo J Ricart Arbona

## Abstract

The soles of staff shoes accessing vivaria can become contaminated on urban streets, potentially serving as a source of fomite-mediated transmission of adventitious agents to laboratory rodents. While shoe covers may mitigate this risk, donning them can lead to hand contamination. Staff accessing our vivaria use motor-driven shoe cleaners hundreds of times daily to remove and collect particulates via a vacuum collection system from the top, sole, and sides of shoes instead of shoe covers. Shoe cleaner debris (SCD) and contact media (CM) exposed to SCD from shoe cleaners in 5 vivaria were assessed by PCR for 84 adventitious agents. SCD and CM samples tested positive for 33 and 37 agents, respectively, and a combined 39 agents total. To assess SCD infectivity, NSG and Swiss outbred mice were housed for 7 days in direct contact with SCD and oronasally inoculated with a suspension created from SCD collected from each of the 5 vivaria. Mice were tested by PCR and serology 3-, 7-, 14- and 63-days post-inoculation. All mice remained healthy until the study’s end and tested negative for all agents found in SCD/CM except murine astrovirus 1, *Staphyloccocus xylosus* and *Candidatus Savagella*, agents known to be enzootic in the experimental mouse source colony. In a follow-up study, the soles of 27 staff street shoes were directly sampled using CM. Half of CM was used for PCR, while the other half was added as bedding material to a cage containing NSG and Swiss outbred mice. While CM tested positive for 11 agents, all mice were healthy 63 days post-exposure and again positive for only enzootic agents. These results suggest that shoe debris might not be a significant biosecurity risk to laboratory mice, questioning the need for shoe covers or cleaners when entering experimental barrier vivaria.

## Introduction

A 2020 survey revealed that over 2 million mice and rats were reported outside of New York City (NYC) buildings over a 90-day period.^36^ While the precise number of wild rodents in NYC is unknown, statistical models estimate that a population of approximately 2 million rats cohabitate with over 8 million city residents.^4^ Wild mice in NYC have been found to carry many infectious agents in their feces including *Shigella* spp.; enteroinvasive, enteropathogenic and Shiga toxin-producing *Escherichia coli*; *Clostridioides difficile*; *Clostridium perfringens*; *Salmonella enterica*; *Leptospira* spp.; murine adenovirus 2; murine parvovirus 2; murine polyomavirus; lactate dehydrogenase-elevating virus; murine hepatitis virus; and murine rotavirus among other agents.^56–57^ Our feral mouse monitoring program has identified many of these agents in mice captured in proximity to our research buildings with vivaria. In fact, we suspect a wild rodent incursion to have been the likely cause of an outbreak of a novel virus, murine astrovirus 2, in one of our vivaria.^46^ A recent New York Times article discussed how the number of wild rodents and the incidence of zoonotic diseases, for which these animals serve as reservoirs, increased in NYC during the COVID 19 pandemic.^6^ This urban wild rodent population is not only a threat to human health, but also poses a significant biosecurity threat to research rodent colonies, as many of the infectious agents they carry are well-documented to confound experimental outcomes in research rodents.^5,7,22^

To mitigate this risk, animal care programs strive to exclude specific agents from their rodent colonies by implementing stringent biosecurity practices and colony health monitoring programs.^5,7,9,13,19,33^ Given the seemingly ubiquitous presence of wild rodents on NYC streets and in subway stations, it is highly probable that research and animal care staff unwittingly accumulate debris shed by wild rodents as well as their excrement on the soles of their shoes during their travels to and from work. Consequently, street shoes are a plausible source of fomite-mediated transmission of excluded infectious agents to laboratory rodents.

At our institutions, shoe cleaners are utilized to mechanically remove particulate debris from research and animal care staffs’ shoes prior to vivarium entry. Shoe cleaners were adopted for vivarium use from the electronics manufacturing industry, where cleanrooms are used to ensure particulates are excluded from specific manufacturing areas as they can interfere with the function of electronic components. Shoe cleaners consist of motor-driven brushes that remove large and medium-sized particles from the top, bottom, and sides of shoes to reduce the potential that particulates carrying infectious agents are introduced into the vivarium. As these particles are removed, they are collected through a vacuum system into a collection unit. Although uncommon in research vivaria, shoe cleaners are utilized in lieu of shoe covers in our vivaria because donning shoe covers may lead to the contamination of personnel’s hands as they enter the vivarium, increasing the chance of the inadvertent introduction of adventitious agents into the vivarium.^2,17,27,28,34,54^ Despite significant concern among human healthcare experts that shoes can serve as fomites, there is no conclusive evidence as to whether footwear does or does not pose a substantial transmission risk to animals or humans.^32,44,58^ We limit shoe cover use to situations when gross contamination of footwear is likely to occur (e.g., large animal holding rooms and necropsy) or within areas where hazardous agents are used to reduce the likelihood of hazardous agent contamination outside of the containment area. For similar reasons, human hospitals no longer recommend the use of shoe covers as a form of personal protective equipment (PPE) to reduce the introduction of infectious agents and only use them to protect from blood and bodily fluid contamination.^28,34,36,54^

At our institutions, shoe cleaners are used upon entry to 5 vivaria by hundreds of staff daily. Their use presented a unique opportunity to evaluate whether contaminated shoes pose a substantive biosecurity risk to rodent colonies. In this study we aimed to survey shoe debris collected by shoe cleaners or directly from shoe soles for the presence of adventitious agents using multiplex PCR and confirm if the material collected can transmit infectious agents to mice. Results from this study provides valuable insight as to whether shoes pose a substantive biosecurity risk associated with staff entering a vivarium.

## Materials and Methods

### Experimental Design

#### Vivarium shoe cleaner study

A study was conducted to assess the potential transmission of murine adventitious agents to mice exposed to debris collected from shoe cleaners. Shoe cleaner debris (SCD) was collected from 5 NYC vivaria (designated A to E; Figure 1). We surveyed for 84 infectious agents present in each SCD sample by analyzing 25mL of unaltered SCD, as well as contact media (CM; PathogenBinder™, Charles River Laboratories, Wilmington, MA) exposed to the same 25mL of SCD, by multiplex PCR (Table 1).

**Figure 1.**
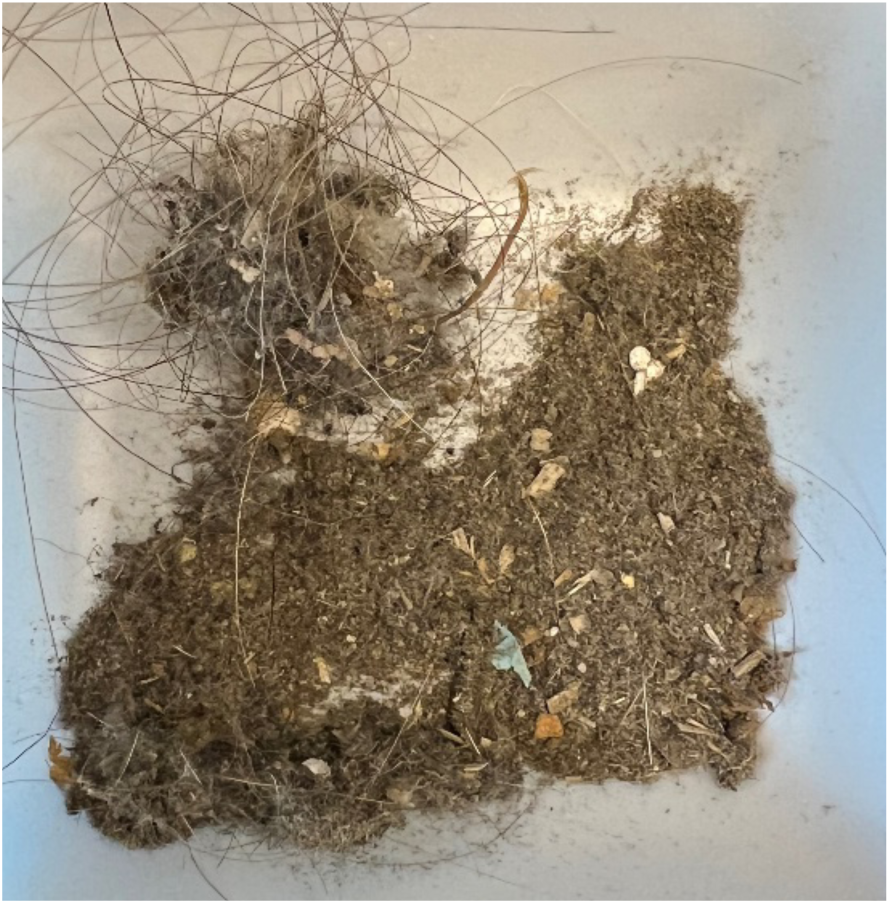
Shoe cleaner debris (SCD) sample without magnification.

**Table 1.**
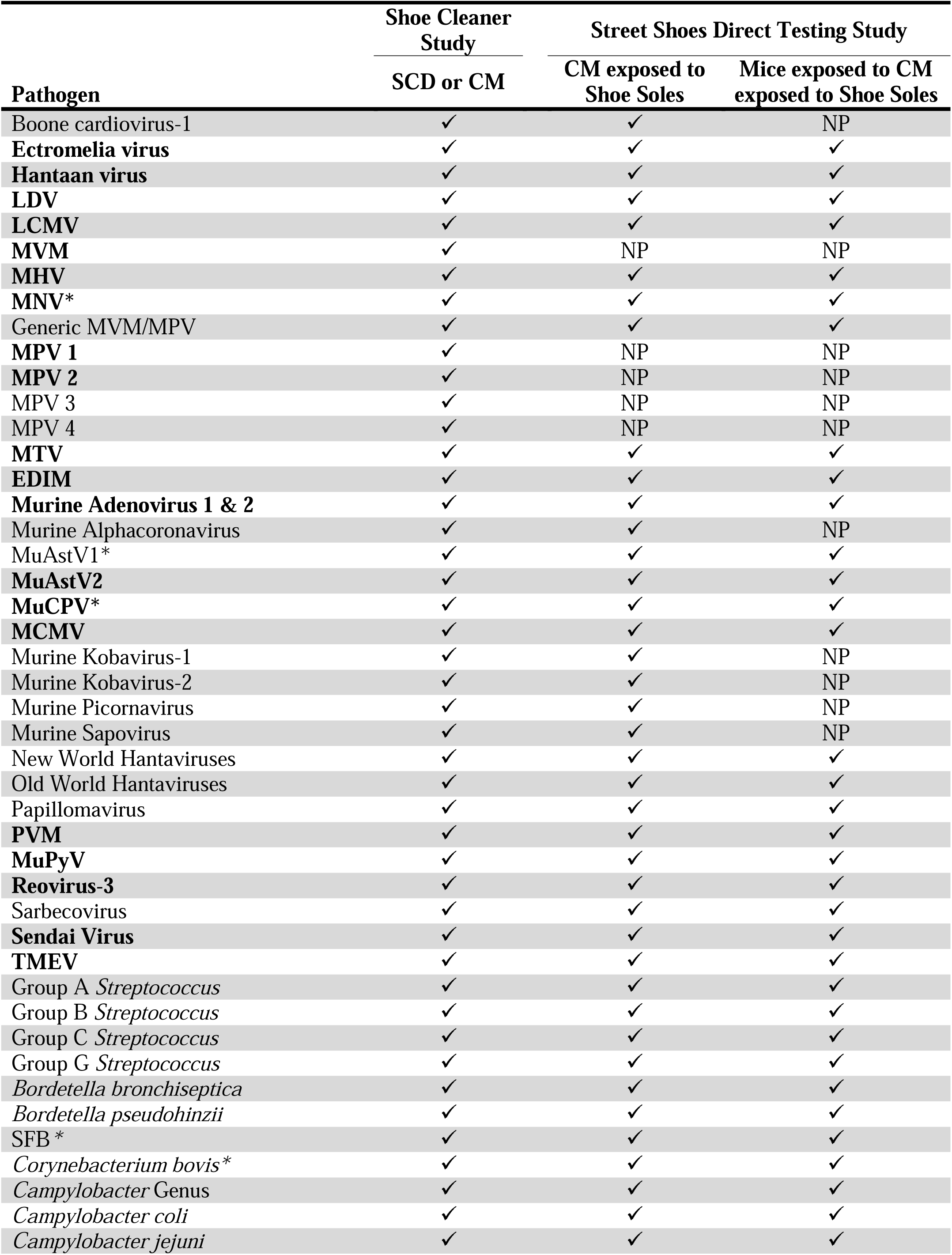
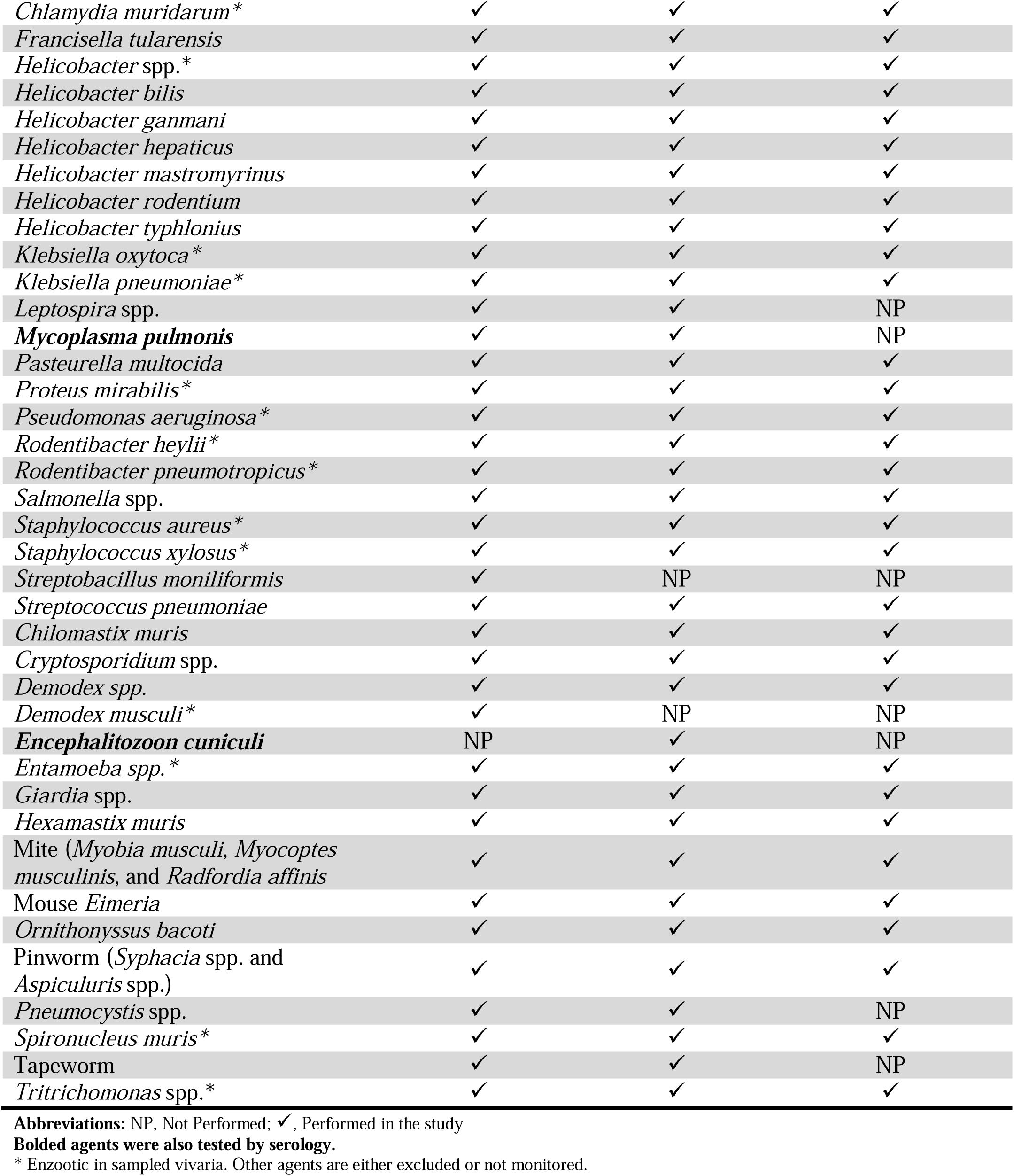
List of Viral, Bacterial and Protozoal Agents Tested by Serology and/or PCR

The SCD from each of the 5 vivaria (5 mice/vivarium) was also used to expose cohoused, naïve, immunodeficient NSG (n=3) and immunocompetent Swiss Webster (SW; n=2) mice. Control mice (n=3 NSG and n=2 SW cohoused in a single cage) were exposed to autoclaved SCD pooled from all 5 vivaria. Mice were inoculated with a SCD suspension orally (PO) and intranasally (IN) as well as by direct contact (DC) for 7 days by adding 25 mL of unaltered SCD to the cage bedding (Figure 2). Following a 7-day exposure period, mice were transferred into a clean autoclaved cage and monitored for up to 63 days post inoculation (DPI). Detailed procedures for collecting and processing of the SCD samples are described in subsequent sections.

**Figure 2.**
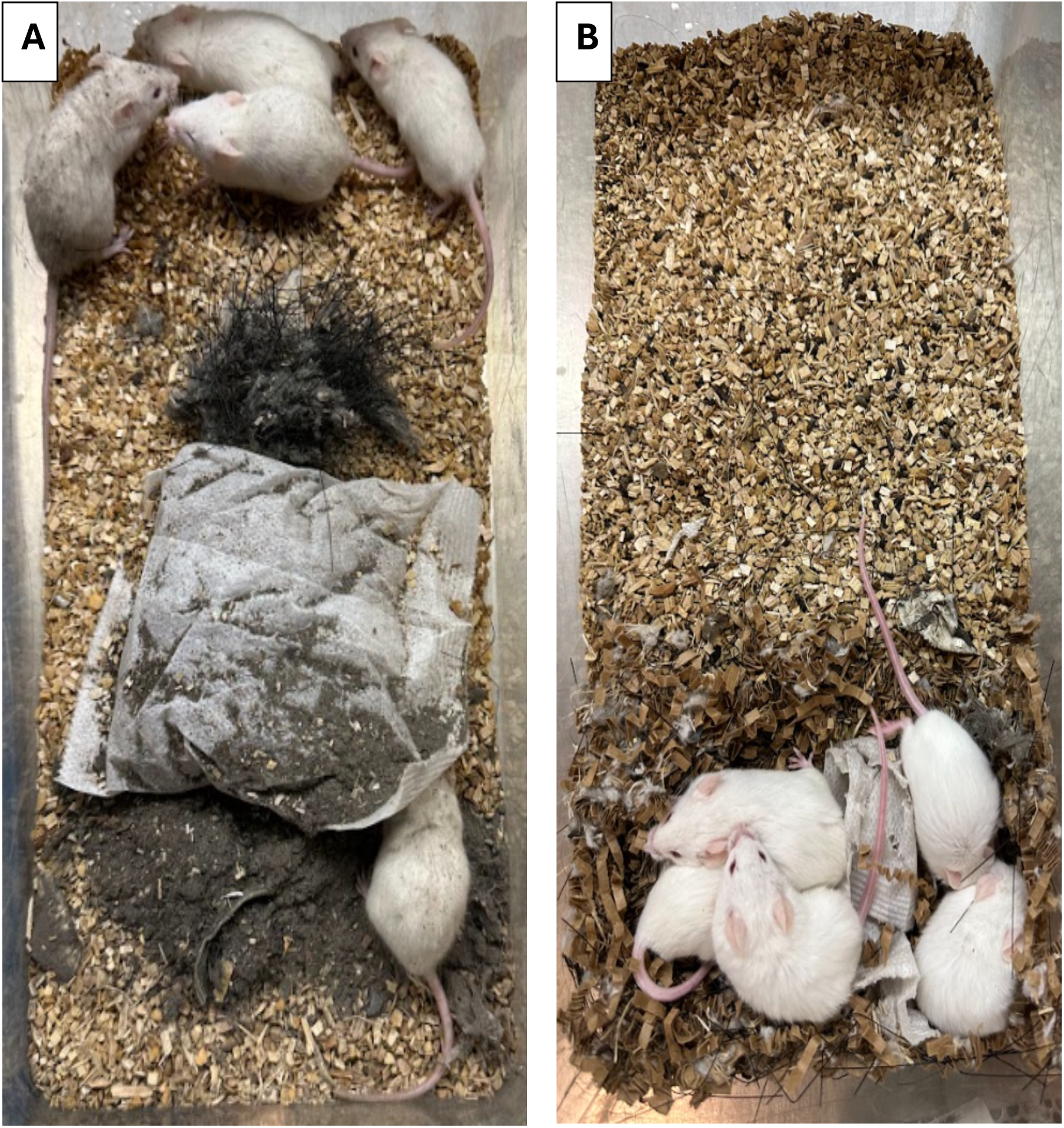
Mice after exposure to shoe cleaner debris (SCD) in the Vivarium shoe cleaner study. (A) Mice following initial exposure to SCD. (B) Mice 1-week post-exposure to SCD with some of the debris integrated into the nest.

Fecal pellets as well as fur and oral swabs were collected from each mouse on 0, 3, 7, 14 and 63 DPI. All samples from 0, 14 and 63 DPI were pooled by day and cage, while samples collected on 3 and 7 DPI were combined into a single sample and pooled by cage. Samples were stored at −80°C (−112°F) until they were tested by PCR for 84 murine and zoonotic viruses, bacteria and parasites. One NSG mouse per cage was euthanized on 7 and 14 DPI while the remaining NSG and 2 SW mice were euthanized on 63 DPI. Terminal blood collection (SW only) was performed immediately after euthanasia and sera was stored at −80°C (−112°F) until tested for 20 agents by multiplexed fluorometric immunoassay (MFIA^®^; Charles River Laboratories). Following euthanasia, deep skin scrapes of the cervical, thoracic and sacral skin and a complete necropsy were performed.

To confirm results from the initial shoe debris exposure, an equal volume of SCD (6 mL/vivarium) were pooled and used to expose 2 cages, each with 5 NSG female mice as described previously. The mice were monitored for clinical signs and euthanized on DPI 63. A gross necropsy was performed on each mouse. No further analyses were performed as mice were healthy and lesion free.

#### Street shoes direct testing study

As SCD had accumulated over a period of up to 335 days and was subject to recognized and unrecognized environmental stressors during collection and storage, an additional study was conducted to detect the presence of infectious agents collected directly from the soles of 27 animal care staff’s shoes on building entry for their assigned shift. The samples were collected on April 26^th^, 2024. The temperature for the preceding 24 hours was 46 – 60°F and there had been a light drizzle (less than <0.1inches of precipitation).^40^ Detailed procedures for collecting and processing of the street shoe samples are described in subsequent sections.

Within 2 hours of swabbing the soles with CM, each CM was divided in half and were divided into 2 groups. One group was added directly to a cage containing 3 NSG and 2 SW female mice while the other group was tested by PCR for 75 infectious agents (Table 1). Following a 7-day exposure period, mice were transferred into a new autoclaved cage and monitored until 63 DPI at which point, they were euthanized and deep skin scrapes of the cervical, thoracic, and sacral skin, and a complete necropsy was performed.

### Animals

Five- to six-week-old female NOD.Cg-Prkdc^scid^ Il2rg^tm1wjl^/SzJ (n = 31; NSG, The Jackson Laboratory, Bar Harbor, ME) and 5- to 6-week-old female Tac:(SW) (Swiss Webster; *n* = 14, Taconic Biosciences, Germantown, NY) mice were utilized in this study. All mice were individually ear-tagged, assigned a distinct numerical identifier, and randomly assigned to appropriate groups (https://www.random.org/lists/).

All NSG and SW mice were free of mouse hepatitis virus (MHV), Sendai virus, mouse parvovirus (MPV), minute virus of mice (MVM), murine norovirus (MNV), murine astrovirus 2 (MuAstV2), pneumonia virus of mice (PVM), Theiler meningoencephalitis virus (TMEV), mouse rotavirus (EDIM), Reovirus type 3, lymphocytic choriomeningitis virus (LCMV), lactate dehydrogenase elevating virus (LDV), K virus, mouse adenovirus 1 and 2 (MAD 1/2), murine polyoma virus (MuPyV), murine cytomegalovirus (MCMV), mouse thymic virus (MTV), Hantaan virus, murine chaphamaparvovirus-1(MuCPV), *Mycoplasma pulmonis*, CAR *bacillus* (*Filobacterium rodentium*), *Chlamydia muridarum*, *Citrobacter rodentium, Rodentibacter pneumotropicus, Helicobacter* spp., *Salmonella* spp., *Streptococcus pneumoniae*, Beta-hemolytic *Streptococcus* spp., *Streptobacillus moniliformis, Chlamydia muridarum, Clostridium piliforme, Corynebacterium bovis, Corynebacterium kutscheri, Staphylococcus aureus, Klebsiella pneumoniae, Klebsiella oxytoca*, *Pseudomonas aeruginosa, Myobia musculi*, *Myocoptes musculinis*, *Radfordia affinis*, *Syphacia* spp. *Aspiculuris* spp., *Demodex musculi*, *Pneumocystis* spp, *Giardia muris*, *Spironucleus muris*, *Entamoeba muris*, *Tritrichomonas muris*, and *Encephalitozoon cuniculi* at the initiation of the studies.

### Husbandry and housing

The study was conducted in a dedicated quarantine facility operated at ABSL-2.^36^ Mice were maintained in sterile individually ventilated polysulfone cages with stainless-steel wire bar lids (WBLs) and filter tops (# 19, Thoren Caging Systems, Hazelton, PA) on aspen chip bedding (PWI Industries, Quebec, Canada) at a density of 5 mice per cage. Each cage was provided with a bag constructed of Glatfelter paper containing 6 g of crinkled paper strips (EnviroPak^®^, WF Fisher and Son, Branchburg, NJ) and a 2-inch square of pulped virgin cotton fiber (Nestlet, Ancare, Bellmore, NY) for enrichment. Each cage (including cage bottom, bedding, WBL and water bottle) was changed weekly. Cages were only opened or changed within a class II, type A2 biological safety cabinet (BSC; LabGard S602-500, Nuaire, Plymouth, MN). All mice were fed a natural ingredient, closed source, autoclavable diet (5KA1, LabDiet, Richmond, VA) and provided ad libitum autoclaved reverse osmosis acidified (pH 2.5 to 2.8 with hydrochloric acid) water in polyphenylsulfone bottles with autoclaved stainless-steel caps and sipper tubes (Techniplast, West Chester, PA). The rooms were maintained on a 12:12-h light:dark cycle (6 AM on:6 PM off), relative humidity of 30% to 70%, and room temperature of 72 ± 2°F (22 ± 1°C).

Caging was autoclaved using a pulsed vacuum cycle of 4 pulses at a maximum pressure of 12.0 psig (6.9 kPa), with sterilization temperature of 121°C (250°F) for 20 min and a 10.0 in Hg vacuum dry (3.4 kPa). Sterilization was confirmed by autoclave tape color change and post hoc verification of cycle chamber operating conditions. In addition, autoclave performance was verified weekly using biologic indicators (Attest Biologic Indicators, 3M, Saint Paul, MN). Water bottles were autoclaved at a temperature of 121°C (250°F) for 45 min with a purge time of 10 min.

The animal care and use program at Memorial Sloan Kettering Cancer Center (MSK) is accredited by AAALAC International and all animals are maintained in accordance with the recommendations provided in the Guide.^29^ All animal use described in this investigation was approved by MSK’s IACUC and conducted in agreement with AALAS position statements on the Humane Care and Use of Laboratory Animals and Alleviating Pain and Distress in Laboratory Animals.

### Sample collection and processing

#### Vivarium shoe cleaner study

SCD was collected from the shoe cleaner collection reservoirs from each of 5 vivaria A through E. Details regarding the number of laboratory staff that have access to each vivaria and its’ average daily cage census are detailed in Table 2 to provide context as to the number of people (animal care staff excluded) utilizing the shoe cleaners. Samples were pooled and thoroughly mixed in vivaria with multiple shoe cleaners (A and C). The shoe cleaning system in vivarium A and C consists of 5 individual shoe cleaners (Shoe Brush 2001-TB, Liberty Industries Inc., East Berlin, CT) each connected through a centralized vacuum system to a reservoir into which the SCD from the associated cleaner is collected and stored. Approximately 100 mL of debris from each of the aforementioned shoe cleaner reservoirs from the same building was pooled and processed. Vivaria B, D, and E each have a single shoe cleaner (Shoe Brush 2010SC, Liberty Industries Inc.) that collects debris directly into a collection bag located within the shoe cleaner. Debris (500mL) from each collection bag was collected and processed. Samples from each vivaria were aliquoted into five 50mL sterile conical tubes (Falcon^®^ 50mL High Clarity PP Conical Bottom Sterile Centrifuge Tube, 352070, Corning Inc., Corning, NY) containing approximately 25mL of SCD. Excess SCD was stored for use as needed in confirmatory studies. Aliquots from each vivaria were allocated for direct PCR testing, exposure of CM which was subsequently tested by PCR, preparing a suspension for intranasal and oral (IN/PO) inoculation of mice, exposing mice by DC, and autoclaving for exposure of mice serving as Controls.

**Table 2.** Cage and User Census per Vivaria

| **Vivarium** | **Number of Laboratory Staff**<br>**with Access to Vivarium** | **Average Daily**<br>**Cage Census** |
| --- | --- | --- |
| **A** | **306** | **11285** |
| **B** | **34** | **1529** |
| **C** | **537** | **25552** |
| **D** | **41** | **1660** |
| **E** | **100** | **6222** |

To prepare the CM for PCR testing, a 25mL SCD aliquot from each vivaria and the autoclaved Control SCD were each transferred into individual autoclaved plastic boxes (PathogenBinder™ kit box, Charles River Laboratories) containing a new CM. The sample was processed according to the manufacturer’s instructions, which included shaking the box 20 times horizontally and 20 times vertically within a BSC (LabGard S602-500, Nuaire) to ensure full exposure of the CM to the SCD over a 15-20 second period.^12^ Precautions were taken to prevent cross-contamination of CM from different vivaria during sample processing including changing gloves and disinfecting all interior surfaces of the BSC between samples with a 0.5% accelerated hydrogen peroxide disinfectant (Peroxigard RTU, Virox Technologies, Ontario, CA) allowing for a minimum of 5 minutes contact time before wiping up the solution with a cloth (WypAll Economizer L30 ¼ Fold Wipers, Kimberly-Clark, Roswell, GA).

To prepare the IN/PO inoculum, a 15mL aliquot from each SCD was transferred to a 50mL sterile conical tube, sterile phosphate buffered saline (15mL) was added, and the tube was vortexed for 60 seconds to ensure homogeneity followed by centrifugation at 10,000 g for 20 minutes. The supernatant was collected and administered by the IN and PO routes within 1 hour of preparation.

Control SCD was prepared by pooling 25mL aliquots of SCD from each vivarium in a microisolator cage (#19, Thoren Caging Systems) with a filter top secured utilizing heat sensitive autoclave tape (Medline, Mundelein, IL). The cage was then placed within an autoclavable biohazard bag (Part No. 848, Autoclavable Biohazard Waste Bags with Polypropylene w/ Indicator, Medegen Medical Products, Hauppauge, NY) and taped shut with heat sensitive autoclave tape. The bagged cage was then autoclaved using the cycle parameters described above. Sterilization was confirmed by autoclave tape color change and post hoc verification of cycle chamber operating conditions. Following autoclaving, control debris was prepared and utilized as described.

For the confirmatory study, 3mL of SCD from each vivarium stored at room temperature for 3 months was pooled with approximately 3mL of freshly collected debris from the same vivaria, to inoculate NSG mice with pooled SCD. NSG mice were inoculated through PO, IN and DC exposure as described above. In total, these cages were inoculated with 30mL of pooled SCD representing all vivaria.

#### Street shoes direct testing study

Samples were collected by firmly pressing a new CM onto the heel of the sole, moving it from the heel to the toe to the heel, and then repeating the process on the other sole. All exposed CMs (n=27) were pooled into a single sterile bottle (500mL Sterile Corning™ Octagonal PET Storage Bottle with 31.7mm Screw Caps, Corning Inc.) until processing within an hour after collection. No identifying information was collected from volunteers.

Each CM was cut in half on the diagonal using autoclaved stainless steel iris scissors. Half of each CM (n=27) were combined and introduced into a single cage housing 3 NSG and 2 SW mice within 3 hours of collection (Figure 3); the mice were exposed to the CM for 7 days until the next cage change at which point mice they were housed in autoclaved cages for the remainder of the study. The remaining CM halves were returned to the 500mL sterile bottle for PCR testing for 75 murine and zoonotic agents (Table 1).

**Figure 3.**
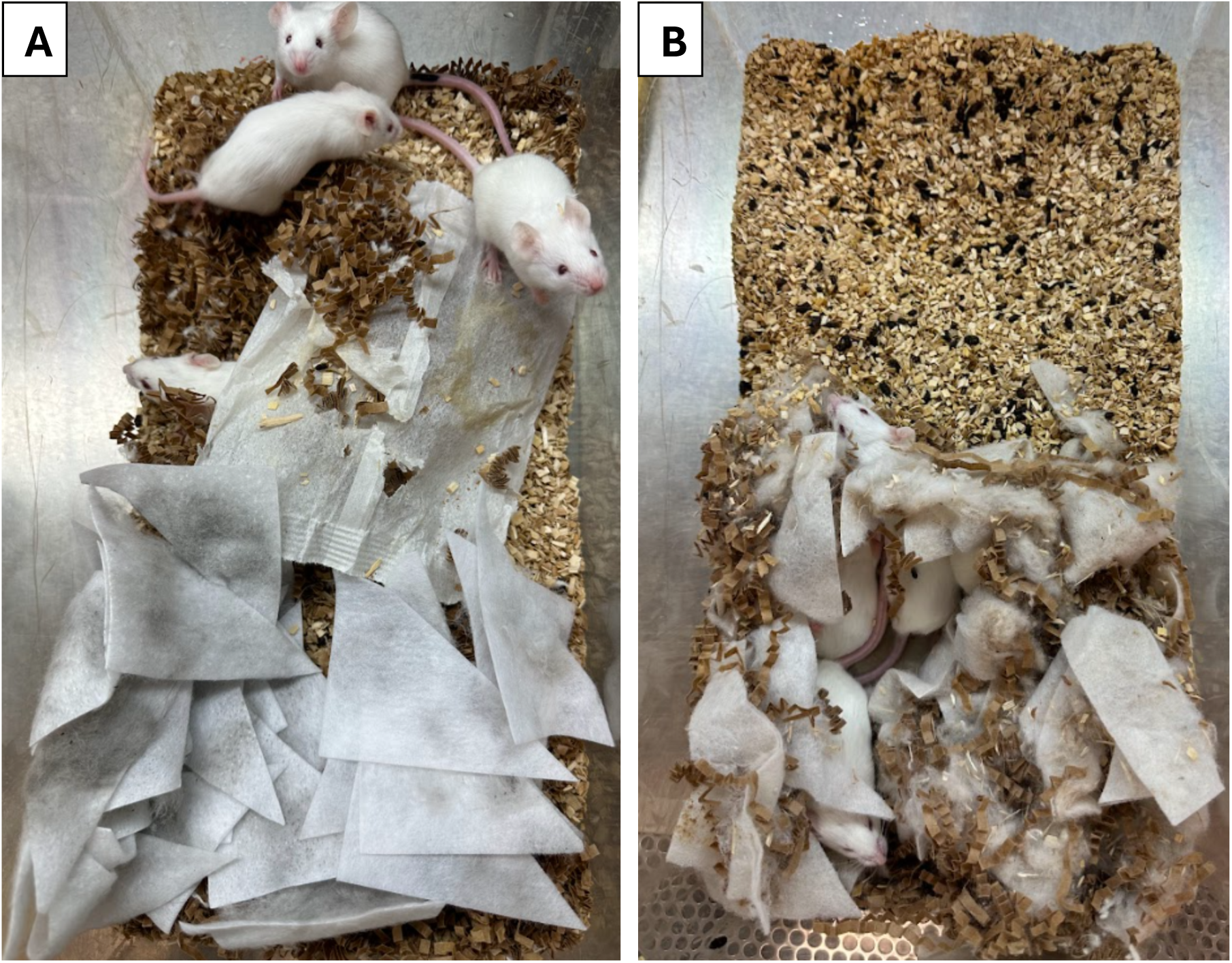
Mice before and after 7 days of exposure to contact media (CM) in the NYC street shoes study. (A) Mice following initial housing with CM. (B) Mice 1 week following CM placement. Mice have created a nest using CM.

### Inoculation with and exposure to shoe cleaner debris

Control mice were exposed prior to the experimental mice. The oral inoculum (0.3 mL) was administered via a 1mL sterile syringe (BD 1mL clear Luer-Lok Tip™ syringe, Becton Dickson, Franklin Lakes, NJ) by gently restraining the mice with a three-finger restraint technique, placing the syringe hub on the animals’ lips, and allowing it to lap up the oral inoculum. Each restrained mouse then received 10 microliters of IN sample into each nares via pipette and held vertically for at least 30 seconds post-administration to limit inoculum loss. DC exposure involved depositing 25mL of SCD from each vivarium onto the bedding surface of the cage housing the mice (Figure 2). Between each experimental cage, a new pair of gloves was donned and the internal BSC surfaces sprayed with disinfectant solution (Peroxigard RTU, Virox Technologies) allowing for a minimum of 5 minutes contact time before wiping up the solution.

### Clinical evaluation, weight assessments and humane endpoint criteria

All mice were observed cage side daily for morbidity, including but not limited to cyanosis, change in body condition, dyspnea, diarrhea, dehydration, hunched posture, lethargy, oral/nasal/ocular discharge, piloerection and sneezing. All mice were weighed prior to study initiation to provide a baseline weight. If clinical signs of disease developed, mice were weighed daily and euthanized if they lost ≥20% of their baseline weight.

### Fecal pellet, fur swab, oral swab and terminal blood collection

Fecal pellets for multiplex PCR analysis were collected directly from each mouse. Mice were lifted by the base of the tail and allowed to grasp the cage wire bar lid while a sterile 1.5mL microfuge tube was held below the anus to allow feces to fall directly into the tube. If the mouse did not defecate within a 30-s period, it was returned to the cage and allowed to rest for at least 2 min before collection was reattempted until a sample was produced. Fur was swabbed using sticky sterile swabs (Pink “Sticky” Swab, Charles River Laboratories), starting at the tail base moving over the dorsum and head finishing at the nose. The ventrum was then swabbed, starting from the caudal abdomen moving cranially ending at the chin. Oral samples were collected using non-alginate swabs (Oral Swab, Charles River Laboratories) by gently restraining the mice with the and sampling the left and right buccal mucosa as well as the dorsal surface of the mouse’s tongue.^31^ Samples were pooled by cage with up to 10 fecal pellets and up to 10 oral and/or fur swabs in a single tube. Terminal blood collection was performed on each SW mouse via intracardiac puncture immediately after euthanasia. Clotted blood was centrifuged and sera was stored at −80°C (−112°F) until testing.

### Euthanasia and pathology

Mice were euthanized by carbon dioxide (CO_2_) asphyxiation in accordance with accepted recommendations.^2^ After euthanasia, a deep skin scrape of cervical, thoracic and sacral skin was performed, and then a complete necropsy was performed on each mouse. Gross lesions were recorded and sterile samples from the mandibular and mesenteric lymph nodes were collected, stored at −80°C (−112°F), and tested by PCR. In the *Vivarium shoe cleaner study,* identifiable lymph nodes were harvested and frozen at -80°C for PCR testing for all agents. Full-thickness skin samples (diameter, 6 mm) were collected by punch biopsy (Integra, Mansfield, MA) or autoclaved stainless steel iris scissors from the pinnae, head, interscapular skin, mid-ventrum, mid-dorsum, caudal dorsum, and anterior aspect of the limbs of mice that were PCR positive for *Demodex*. Sections of heart, thymus, lungs, liver, gallbladder, kidneys, pancreas, stomach, duodenum, jejunum, ileum, cecum, colon, lymph nodes (mandibular, mesenteric), salivary glands, skin (trunk and head), urinary bladder, uterus, cervix, vagina, ovaries, oviducts, adrenal glands, spleen, thyroid gland, esophagus, trachea, spinal cord, vertebrae, sternum, femur, tibia, stifle joint, skeletal muscle, nerves, skull, nasal cavity, oral cavity, teeth, ears, eyes, pituitary gland, and brain were collected and fixed in 10% neutral buffered formalin for at least 72 hrs. After formalin fixation, the skull, spinal column, sternum, femur, and tibia were decalcified in a formic acid and formaldehyde solution (Surgipath Decalcifier I, Leica Biosystems, Wetzlar, Germany). All fixed tissues were then processed in ethanol and xylene in a tissue processor and embedded in paraffin (Leica ASP6025, Leica Biosystems). Paraffin blocks were sectioned at 5-μm and stained with hematoxylin and eosin (H&E). All tissues were examined by board-certified veterinary pathologists (MA and SC).

### Immunohistochemistry (IHC)

Selected sections of large intestine from SW mice were stained for Chlamydia major outer membrane protein (MOMP). Briefly, formalin-fixed paraffin-embedded sections were stained using an automated staining platform (Leica Bond RX, Leica Biosystems). Following deparaffinization and heat-induced epitope retrieval in a citrate buffer at pH 6.0, the primary antibody against Chlamydia MOMP (NB100-65054, Novus Biologicals, Centennial, CO) was applied at a dilution of 1:500. A rabbit anti-goat secondary antibody (Cat. No. BA-5000, Vector Laboratories, Burlingame, CA) and a polymer detection system (DS9800, Novocastra Bond Polymer Refine Detection, Leica Biosystems) was then applied to the tissues. The 3,3’-diaminobenzidine tetrachloride (DAB) was used as the chromogen, and the sections were counterstained with hematoxylin and examined by light microscopy. Intestines from NSG mice experimentally infected with *Chlamydia muridarum* were used as positive control.^51^

### PCR and multiplexed fluorometric immunoassay (MFIA)

Infectious agent nucleic acid was isolated from lysate prepared from samples and PCR testing was conducted using validated, proprietary, real-time fluorogenic 5′ nuclease PCR assays that target rodent pathogens listed in Table 1.^26–27^ If initial testing was positive, DNA was re-isolated from a retained lysate sample which was retested by PCR to confirm the finding. A positive result was reported when the repeated assay was positive based on real-time PCR cycle threshold values equivalent to or greater than established individual assay cutoff(s). To monitor for successful DNA recovery after extraction and to determine whether there may be evidence of PCR inhibitors, a nucleic acid recovery control assay was also performed. Exogenous algae DNA was added to the sample lysate prior to nucleic acid isolation and was subsequently monitored by using a real-time PCR assay targeting the algae sequence. Assays for which nucleic acid recovery control assays for samples had greater than a log_10_ loss of template copies compared with control wells were not accepted as valid.

Target nucleic acid copies per reaction in a sample were estimated by comparing the cycle threshold (Ct) values of the average sample and 100-copy positive template control; a difference of 3.3 Ct corresponds to approximately a 10-fold difference in copy number.^52^ This method was used to provide an estimated target template copy number per reaction to assess the positive signal for samples and differences in values across groups. This method was not considered to be quantitative PCR, which requires triplicate replicates of control template dilutions and sample nucleic acid, therefore relative copy numbers were reported.

For several agents (parvoviruses, *Campylobacter* spp., *Helicobacter* spp., and *Demodex* spp.) prescreening was conducted to determine if testing by species was necessary. In cases in which the aforementioned assays were positive, species testing was performed. In these cases, the prescreening agent was not included in the total agent count; only the positive species level results were counted.

A MFIA for MuAstV-2, Ectromelia, MAD 1/2, MHV, MNV, MPV-1, MPV-2, MVM, LDV, LCMV, MCMV, MTV, EDIM, MuPyV, PVM, Reovirus-3, Sendai Virus, TMEV, Haantan virus, *Mycoplasma pulmonis,* and *Encephalitozoon cuniculi* was conducted on sera using an established validated MFIA platform.^60^ For each assay, the net median fluorescence intensity signal was calculated by subtracting the tissue control from the antigen median fluorescence intensity. Net scores were calculated from net MFIA values based on cutoffs and formulas as described in the manual. Values of <1.5 and ≥3 were classified as negative and positive, respectively; net signals between these cutoffs were identified as equivocal/indeterminate.^11^

### Statistical Analysis

Odds ratios (ORs) were calculated for each of the 6 vivaria to estimate the strength of association between positive SCD PCR and positive PCR samples in CM exposed to SCD. Chi-square analysis was used to evaluate whether statistically significance was seen. Descriptive statistics and statistical hypothesis testing were performed using JMP Pro 17 software (SAS Institute Inc., Cary, NC). A P value of less than or equal to 0.05 denoted statistical significance.

## Results

### Vivarium shoe cleaner study

Nucleic acid from 13 viruses, 18 bacteria, and 8 ectoparasites or protozoa were detected by PCR in either or both SCD and CM exposed to SCD; 17 of these agents are enzootic in the colonies housed in the vivaria for which the shoe cleaners are used (Tables 3, 4 and 5). Nucleic acid from 33 agents was detected in SCD, 2 of which, MuAstV2 and MVM, were not detected in the CM. Conversely, nucleic acid from 37 agents was detected in CM, 6 of which, Group B *Streptococcus, Helicobacter hepaticus*, MHV, MPV-3, MuCPV, and rodent papillomavirus were not detected in the SCD. Agents which were identified in SCD and CM 100% of the time included: MuAstV1, *Chlamydia muridarum, Helicobacter typhlonius, Klebsiella oxytoca, Staphylococcus aureus* and *Tritrichomonas* spp. Agents that were identified in SCD or Cm more than 50% of the time included: MAV 1 & 2, MNV, mouse parvoviruses (MVM/MPV), *Corynebacterium bovis, Campylobacter* spp., *Helicobacter ganmani, Helicobacter mastromyrinus, Klebsiella pneumoniae, Proteus mirabilis, Pseudomonas aeruginosa, Staphylococcus xylosus, Demodex* spp., *Entamoeba* spp., *Eimeria*, and *Spironucleus muris.* Agents that were never detected by PCR and/or serology are listed in Supplemental Table 1.

**Table 3.** Positive Viral PCR Results of Shoe Cleaner Debris (SCD) and Contact Media (CM) Exposed to SCD^a^

| Virus | Control SCD | Control CM | Source |  |  |  |  |  |  |  |  |  | %<sup>b</sup> | % Positive | SCD:CM Mismatches |
| --- | --- | --- | --- | --- | --- | --- | --- | --- | --- | --- | --- | --- | --- | --- | --- |
|  |  |  | A SCD | A CM | B SCD | B CM | C SCD | C CM | D SCD | D CM | E SCD | E CM |  |  |  |
| MuAstV1* | + | + | + | + | + | + | + | + | + | + | + | + | 100 | 100.0 | 0 |
| MuAstV2 | - | - | - | - | + | - | - | - | - | - | - | - | 20 | 8.3 | 1 |
| Ectromelia | - | - | - | - | - | - | - | - | - | - | + | + | 20 | 16.7 | 0 |
| MAV 1 & 2 | + | - | + | + | + | + | - | - | - | - | + | - | 60 | 50.0 | 2 |
| MHV | - | - | - | - | - | - | - | - | - | - | - | + | 20 | 8.3 | 1 |
| MNV* | + | + | + | + | + | + | - | + | - | - | - | - | 60 | 58.3 | 1 |
| Generic MVM/MPV | + | - | + | + | + | + | - | - | + | + | + | + | 80 | 75.0 | 1 |
| MPV 1 | - | NP | I | - | + | + | NP | NP | - | - | - | - | 20 | 16.7 | 0 |
| MPV 2 | - | NP | I | - | + | + | NP | NP | - | - | - | - | 20 | 16.7 | 0 |
| MPV 3 | - | NP | I | - | - | - | NP | NP | - | + | - | - | 20 | 8.3 | 1 |
| MPV 4 | - | NP | I | - | - | - | NP | NP | - | - | - | - | 0 | 0 | 0 |
| MuCPV* | - | - | - | - | - | - | - | - | - | + | - | + | 40 | 16.7 | 2 |
| MVM | + | NP | I | - | + | - | NP | NP | - | - | + | - | 40 | 16.7 | 2 |
| Papillomavirus | - | - | - | - | - | - | - | - | - | + | - | - | 20 | 8.3 | 1 |
| Sarbecovirus | - | - | + | - | + | - | + | - | - | - | - | + | 80 | 33.3 | 4 |

**Abbreviations:** +, positive; -, negative; I, Inconclusive; MAV, murine adenovirus; MHV, mouse hepatitis virus; MNV, murine norovirus; MPV, mouse parvovirus; MVM, minute virus of mice; MuCPV, murine chaphamaparvovirus-1; N/A, not applicable; NP, not performed
<sup>a</sup> Agents listed in Table 1 were negative by PCR unless otherwise noted in this table. Refer to Supplemental Table 1 for the list of negative agents
<sup>b</sup> Percent of all Vivaria Positive = # Positive Samples by SCD or CM per Vivaria / # of Vivaria
\* Enzootic in sampled vivaria. Other agents are either excluded or not monitored.
**Green Colored Text = Discrepancy with positive SCD but negative CM**
**Red Colored Text = Discrepancy with positive CM but negative SCD**

**Table 4.**
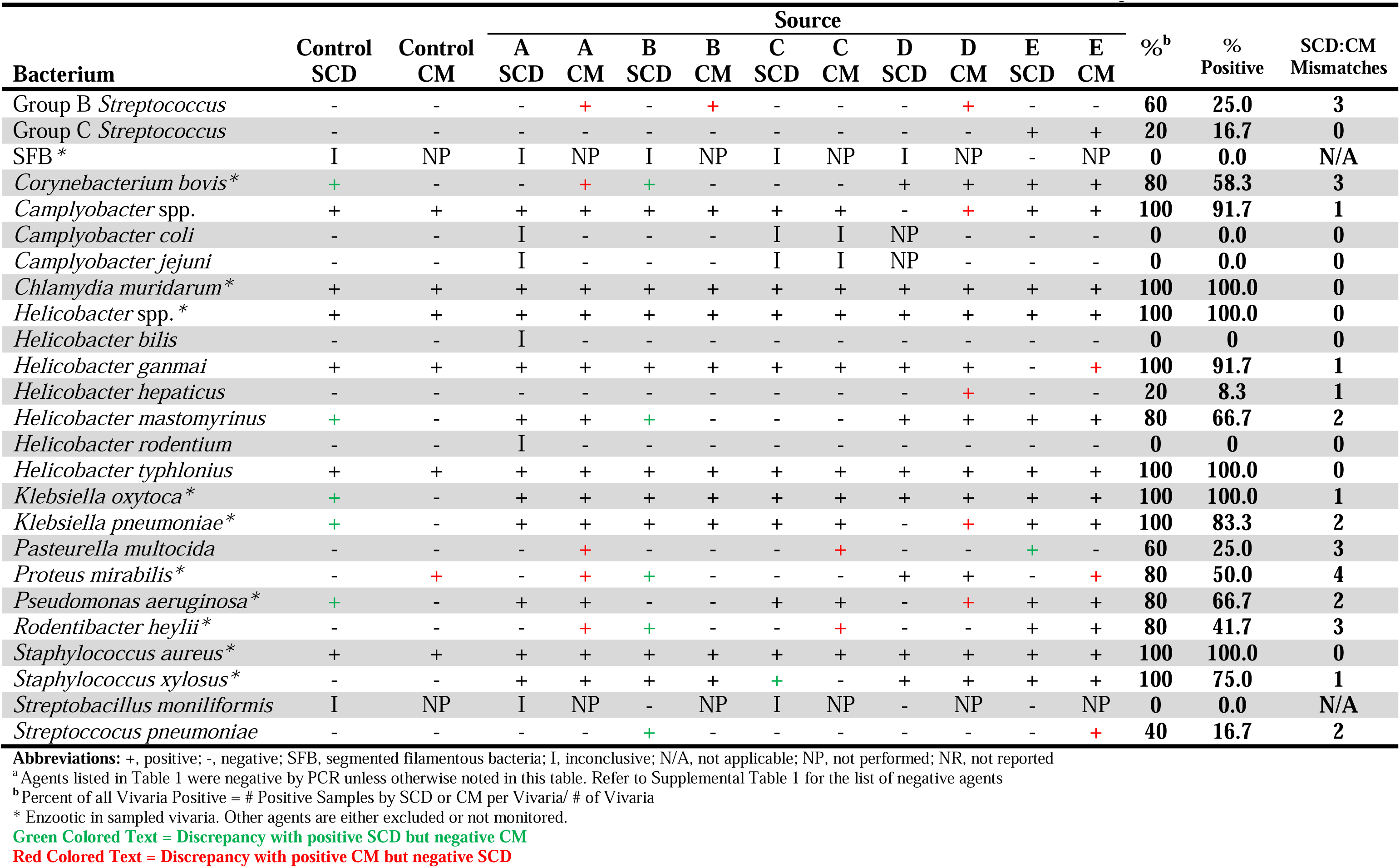
Positive Bacterial PCR Panel Results of Shoe Cleaner Debris (SCD) and Contact Media (CM) Exposed to SCD^a^

| Bacterium | Control SCD | Control CM | Source |  |  |  |  |  |  |  |  |  | %<sup>b</sup> | % Positive | SCD:CM Mismatches |
| --- | --- | --- | --- | --- | --- | --- | --- | --- | --- | --- | --- | --- | --- | --- | --- |
|  |  |  | A SCD | A CM | B SCD | B CM | C SCD | C CM | D SCD | D CM | E SCD | E CM |  |  |  |
| Group B *Streptococcus* | - | - | - | + | - | + | - | - | - | + | - | - | 60 | 25.0 | 3 |
| Group C *Streptococcus* | - | - | - | - | - | - | - | - | - | - | + | + | 20 | 16.7 | 0 |
| SFB* | I | NP | I | NP | I | NP | I | NP | I | NP | - | NP | 0 | 0.0 | N/A |
| *Corynebacterium bovis* * | + | - | - | + | + | - | - | - | + | + | + | + | 80 | 58.3 | 3 |
| *Camphylobacter* spp. | + | + | + | + | + | + | + | + | - | + | + | + | 100 | 91.7 | 1 |
| *Camphylobacter coli* | - | - | I | - | - | - | I | I | NP | - | - | - | 0 | 0.0 | 0 |
| *Camphylobacter jejuni* | - | - | I | - | - | - | I | I | NP | - | - | - | 0 | 0.0 | 0 |
| *Chlamydia muridarum* * | + | + | + | + | + | + | + | + | + | + | + | + | 100 | 100.0 | 0 |
| *Helicobacter* spp.* | + | + | + | + | + | + | + | + | + | + | + | + | 100 | 100.0 | 0 |
| *Helicobacter bilis* | - | - | I | - | - | - | - | - | - | - | - | - | 0 | 0 | 0 |
| *Helicobacter ganmai* | + | + | + | + | + | + | + | + | + | + | - | + | 100 | 91.7 | 1 |
| *Helicobacter hepaticus* | - | - | - | - | - | - | - | - | - | + | - | - | 20 | 8.3 | 1 |
| *Helicobacter mastomyrinus* | + | - | + | + | + | - | - | - | + | + | + | + | 80 | 66.7 | 2 |
| *Helicobacter rodentium* | - | - | I | - | - | - | - | - | - | - | - | - | 0 | 0 | 0 |
| *Helicobacter typhlonius* | + | + | + | + | + | + | + | + | + | + | + | + | 100 | 100.0 | 0 |
| *Klebsiella oxytoca* * | + | - | + | + | + | + | + | + | + | + | + | + | 100 | 100.0 | 1 |
| *Klebsiella pneumoniae* * | + | - | + | + | + | + | + | + | - | + | + | + | 100 | 83.3 | 2 |
| *Pasteurella multocida* | - | - | - | + | - | - | - | + | - | - | + | - | 60 | 25.0 | 3 |
| *Proteus mirabilis* * | - | + | - | + | + | - | - | - | + | + | - | + | 80 | 50.0 | 4 |
| *Pseudomonas aeruginosa* * | + | - | + | + | - | - | + | + | - | + | + | + | 80 | 66.7 | 2 |
| *Rodentibacter heyltii* * | - | - | - | + | + | - | - | + | - | - | + | + | 80 | 41.7 | 3 |
| *Staphylococcus aureus* * | + | + | + | + | + | + | + | + | + | + | + | + | 100 | 100.0 | 0 |
| *Staphylococcus xylosus* * | - | - | + | + | + | + | + | - | + | + | + | + | 100 | 75.0 | 1 |
| *Streptobacillus moniliformis* | I | NP | I | NP | - | NP | I | NP | - | NP | - | NP | 0 | 0.0 | N/A |
| *Streptococcus pneumoniae* | - | - | - | - | + | - | - | - | - | - | - | + | 40 | 16.7 | 2 |

**Abbreviations:** +, positive; -, negative; SFB, segmented filamentous bacteria; I, inconclusive; N/A, not applicable; NP, not performed; NR, not reported
<sup>a</sup> Agents listed in Table 1 were negative by PCR unless otherwise noted in this table. Refer to Supplemental Table 1 for the list of negative agents
<sup>b</sup> Percent of all Vivaria Positive = # Positive Samples by SCD or CM per Vivaria/ # of Vivaria
\* Enzootic in sampled vivaria. Other agents are either excluded or not monitored.
**Green Colored Text = Discrepancy with positive SCD but negative CM**
**Red Colored Text = Discrepancy with positive CM but negative SCD**

**Table 5.**
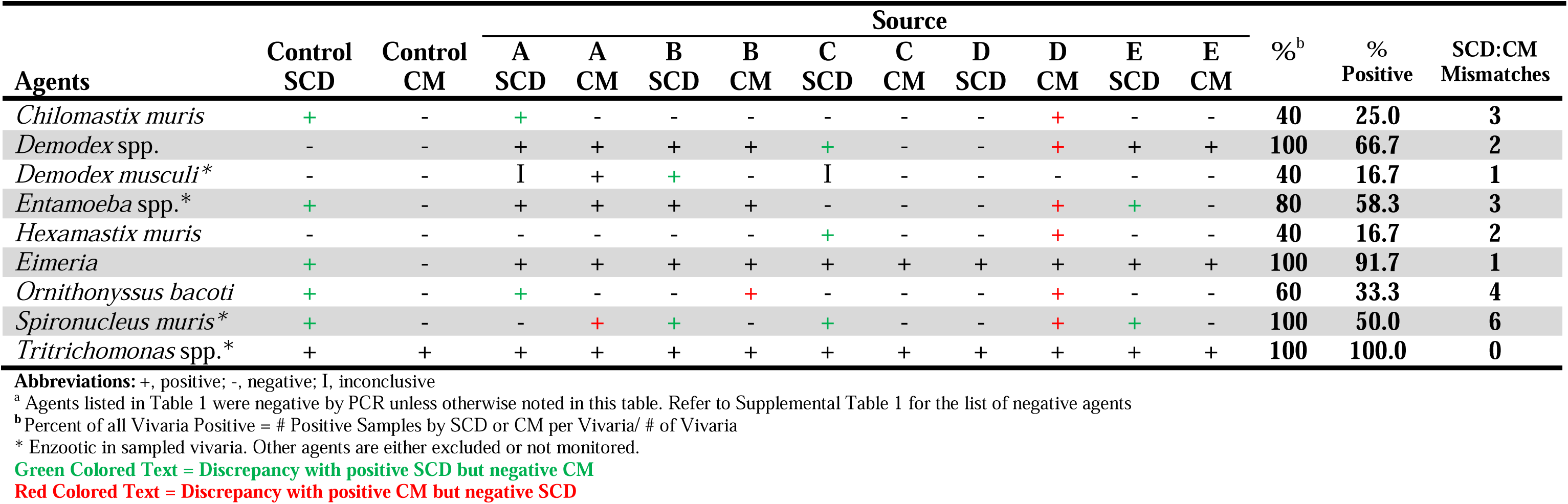
Positive Ectoparasite, Protozoal, and Fungal PCR Results of Shoe Cleaner Debris (SCD) and Contact Media (CM) Exposed to SCD^a^

CM exposed to SCD from Vivarium D was 1.7 times more likely to detect nucleic acid compared to the SCD (P=0.0067). For this vivarium, CM was positive for 28 agents while SCD only detected 14 agents. There were no significant differences in nucleic acid detection between the SCD or CM samples, from vivaria A (OR: 0.79; P = 0.81), B (OR: 1.82; P = 0.22), C (OR: 1.23; P = 0.82), or E (OR: 0.91; P > 0.99). In contrast, autoclaved Control SCD was 2.2 times more likely than CM exposed to autoclaved SCD to detect nucleic acid (P=0.0285). Nucleic acid for 22 agents was detected in the autoclaved control SCD, whereas nucleic acid for only 10 agents was detected in CM. Agents that were identified in 100% of the vivaria, regardless of whether the agent was detected in SCD or CM, included: MuAstV1, *Chlamydia muridarum, Helicobacter ganmani, Helicobacter typhlonius, Klebsiella oxytoca, Klebsiella pneumoniae, Staphylococcus aureus, Staphylococcus xylosus, Demodex* spp., *Eimeria* spp*., Spironucleus muris* and *Tritrichomonas* spp. (Tables 3, 4 and 5). Agents that were identified in 50% or more of the vivaria, regardless of whether the agent was detected in SCD or CM, included: MAV 1 & 2, MNV, mouse parvovirus (MVM/MPV), sarbecovirus, Group B *Streptococcus*, *Corynebacterium bovis, Helicobacter mastomyrinus*, *Pasteurella multocida, Proteus mirabilis, Pseudomonas aeruginosa, Rodentibacter heylii, Entamoeba* spp. and *Ornithonyssus bacoti* (Tables 3, 4 and 5).

SCD and CM results were always in agreement (whether both positive or negative) for 10 agents: MuAstV1, Ectromelia, MPV-1, MPV-2, Group C *Streptococcus*, *Chlamydia muridarum*, *Helicobacter* spp., *Helicobacter typhlonius, Staphylococcus aureus*, and *Tritrichomonas* spp. MuAstV2, *Chlamydia muridarum, Helicobacter spp., Helicobacter typhlonius, Staphylococcus aureus* and *Tritrichomonas* spp. were positive in both SCD and CM 100% of the time. However, discrepancies were noted in the results among samples from the same vivaria for many agents (Table 3, Table 4, and Table 5).

Despite detecting nucleic acid for numerous infectious agents in shoe debris, colonization was not detected by PCR or histopathology, except for potentially *P. aeruginosa.* All mice were positive by PCR for MuAstV1 and *Candidatus Savagella* (SFB), and *Staphylococcus xylosus* prior to exposure (day 0) to SCD, except for mice in group E who were only positive for MuAstV1 and SFB (Table 6), as these agents are not excluded from the vendors’ colonies. Groups remained positive for MuAstV1 and SFB until the last timepoint (Table 6). PCR results for *Staphylococcus xylosus* were temporally inconsistent. While 83% were positive for *S. xylosus* prior to SCD exposure and up to day 7, 100% and 50% were negative on days 14 and 63, respectively. Mice exposed to SCD from Vivarium B tested positive for *Hexamastix muris* and Group B *Streptococcus* on days 3/7 (pooled) only. However, the copy number for both agents was very low with a copy number of 3, and *Hexamastix muris* was negative in both the CM and SCD by PCR (Table 6). Mice exposed to autoclaved SCD and SCD from Vivarium D were positive for *Pseudomonas aeruginosa* on day 63 only. Mice exposed to autoclaved SCD were positive for *Demodex musculi* on day 63 only; however, deep skin scrapes from all mice were negative on day 63. Skin biopsies performed on mice exposed to autoclaved SCD were also negative for *Demodex musculi*. Negative PCR and serology results are provided in Supplemental Table 1.

**Table 6.** Positive Viral, Bacterial, Protozoal and Fungal PCR Copy Number Results of SW/NSG Mice Exposed to Shoe Cleaner Debris (SCD)^a^

| Pathogen | Source |  |  |  |  |  |
| --- | --- | --- | --- | --- | --- | --- |
|  | Control | A | B | C | D | E |
| **Day 0 (unexposed)** |  |  |  |  |  |  |
| **MuAstV1** | 56,960,717 | 28,349,483 | 28,349,483 | 28,349,483 | 56,960,717 | 56,960,717 |
| *Staphylococcus xylosus* | 402 | 50 | 100 | 100 | 1622 | - |
| SFB | 1622 | 402 | 402 | 402 | 1622 | 1622 |
| **Day 3/7** |  |  |  |  |  |  |
| MuAstV1 | 14,109,605 | 7,022,383 | 28,349,483 | 28,349,483 | 56,960,717 | 56,960,717 |
| *Staphylococcus xylosus* | 3,260 | - | 3,260 | 6,549 | 402 | 200 |
| SFB | 807 | 807 | 3,260 | 1,622 | 6,549 | 100 |
| Beta *Streptococcus* Group B | - | - | 3 | - | - | - |
| *Hexamastix muris* | - | - | 3 | - | - | - |
| **Day 14<sup>b</sup>** |  |  |  |  |  |  |
| MuAstV1 | 7,022,383 | 14,109,605 | 14,109,605 | 28,349,483 | 28,349,483 | I |
| *Staphylococcus xylosus* | - | - | - | - | - | - |
| SFB | 1,622 | 402 | 3,260 | 13,159 | 1,622 | I |
| **Day 63** |  |  |  |  |  |  |
| **MuAstV1** | 56,960,717 | 14,109,605 | 14,109,605 | 7,022,383 | 7,022,383 | 430,887 |
| *Staphylococcus xylosus* | 200 | - | - | - | 13,159 | 100 |
| SFB | 214,453 | 3,260 | 6,549 | 3,260 | 402 | 200 |
| *Pseudomonas aeruginosa* | 25 | - | - | - | 1,622 | - |
| *Demodex* spp. | 100<sup>c</sup> | - | - | - | - | - |
| *Demodex musculi* | 12<sup>c</sup> | NP | NP | NP | NP | NP |

**Abbreviations:** -, negative; I, inconclusive; NP, not performed
**Bolded agents had serology performed.** Agents listed in Table 1 were negative by serology except for MuAstV1.
<sup>a</sup>Agents listed in Table 1 were negative by PCR unless otherwise noted in this table. Refer to Supplemental Table 1 for the list of negative agents
<sup>b</sup>Panel inconclusive for the following additional agents in group E on day 14: new world hantavirus, Sendai virus, *Corynebacterium bovis*, *Proteus mirabilis*, *Ornithonyssus bacoti*
<sup>c</sup>Mice negative for *Demodex* spp. via deep skin scrapes of the cervical, thoracic and sacral skin and histologic analysis of head, intrascapular, dorsum, and ventrum biopsies.

Clinical signs were not identified in any mice throughout the study. There was no evidence of pathology associated with infectious agents noted during histologic evaluation of tissues at any timepoint. Incidental or age-associated lesions were observed in a few NSG mice (e.g., extramedullary hematopoiesis, n=6; osseous metaplasia, n=1; and, liver lobe torsion, n=1). While mild to moderate gut-associated lymphoid tissue (GALT) hyperplasia was seen in 5 SW mice euthanized on day 63, MOMP IHC for *C. muridarum* was negative. Mild lymphoid hyperplasia was identified in one or more lymph nodes from 5 SW mice exposed to SCD from Vivarium C, D, E and Control SCD on day 63; however, pooled lymph node biopsies were negative for all agents by PCR. No clinical abnormalities were observed in any of the 10 NSG mice used to confirm results from the initial SCD exposure. No gross lesions were detected in any of the mice necropsied on DPI 63.

### Street shoes direct testing study

In total, nucleic acid for 11 viruses, bacteria, ectoparasites and protozoa were identified on the soles of street shoes (Table 7). Nucleic acid from all these agents, except for *Campylobacter coli*, were also found in the shoe debris. None of the mice exposed to the CM developed clinical signs during the course of the study. Skin scrapings were negative and gross lesions were not detected at necropsy. PCR of feces and MFIA performed on sera from the SW mice were negative for all agents except for MuAstV1 and SFB, the latter were only detected by PCR; however, these agents were present in the mice prior to exposure to CM.

**Table 7.**
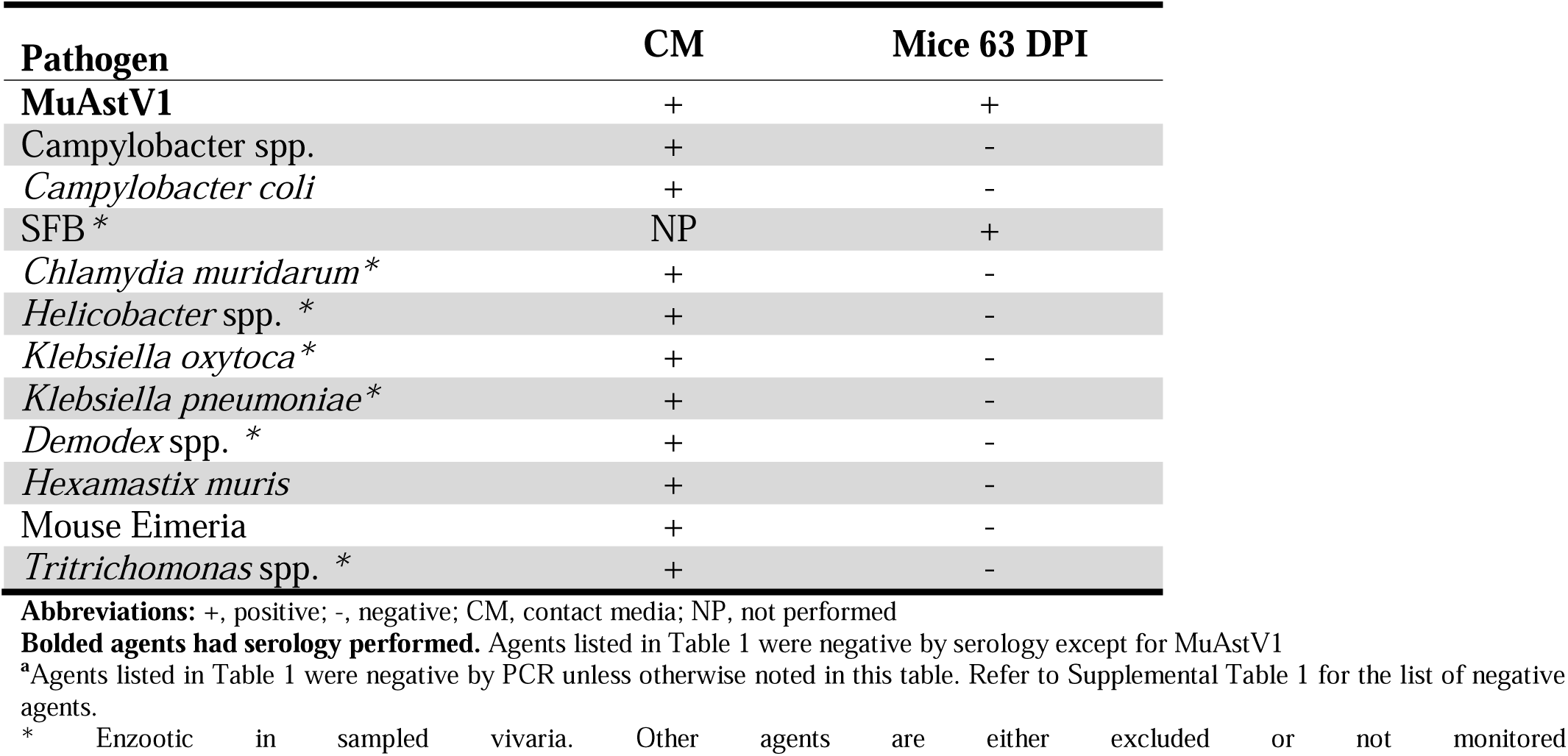
Street Shoes Direct Testing Study PCR Positive Results^a^

| **Pathogen** | **CM** | **Mice 63 DPI** |
| --- | --- | --- |
| **MuAstV1** | + | + |
| **Campylobacter spp.** | + | - |
| *Campylobacter coli* | + | - |
| **SFB\*** | NP | + |
| *Chlamydia muridarum* * | + | - |
| *Helicobacter* spp. * | + | - |
| *Klebsiella oxytoca* * | + | - |
| *Klebsiella pneumoniae* * | + | - |
| *Demodex* spp. * | + | - |
| *Hexamastix muris* | + | - |
| Mouse Eimeria | + | - |
| *Tritrichomonas* spp. * | + | - |

**Abbreviations:** +, positive; -, negative; CM, contact media; NP, not performed
**Bolded agents had serology performed.** Agents listed in Table 1 were negative by serology except for MuAstV1
<sup>a</sup>Agents listed in Table 1 were negative by PCR unless otherwise noted in this table. Refer to Supplemental Table 1 for the list of negative agents.
\* Enzootic in sampled vivaria. Other agents are either excluded or not monitored

## Discussion

Animal care programs strive to exclude specific infectious agents from rodent colonies by implementing stringent biosecurity practices and verify their absence by implementing colony health monitoring programs.^5,33,45^ At our institutions, we utilize mechanical shoe cleaners to remove debris from shoes prior to entering vivaria. Shoe cleaners replaced the use of shoe covers as the latter have questionable effectiveness at preventing entry and spread of potential contaminants and may, in fact, increase the likelihood for contamination of personnel hands and gowns during their donning.^27^ Despite widespread use of shoe covers, and in our case mechanical shoe cleaners, to prevent entry of adventitious agents to vivaria, the risk of shoes serving as fomites for spread of an/or introduction of murine infectious agents has never been directly assessed.^17,27,44^

Our widespread programmatic use of shoe cleaners in an urban environment known to host large numbers of wild mice and rats provided a unique opportunity to assess the biosecurity risk posed by footwear. These wild rodents have been shown to carry MHV, MAV 1 & 2, MNV, MuAstV2, MuCPV, sarbecovirus*, Klebsiella pneumoniae, Campylobacter jejuni, Streptobacillus moniliformis, Cryptosporidium parvum,* and *Ornithonyssus bacoti* among a host of other infectious agents.^18,20,53,56,57^, While we do not exclude all of these agents from our colonies, many are bonafide biosecurity risks. In addition to assessing shoe debris collected by shoe cleaners by PCR for many murine infectious agents, we also sampled the soles of street shoes immediately on entry to the facility and we exposed both immunocompromised and immunocompetent mice to samples to ascertain whether any of these materials could transmit infectious agents to mice.

The study also provided the opportunity to compare the effectiveness of directly testing SCD with the use of CM to indirectly assess the presence of nucleic acid by PCR. While results from SCD and CM were not always congruent, overall, both samples identified a similar number of infectious agents. Discrepancies between both positive and negative results for the same sample and agent occurred 68 times. Since each CM was exposed to the same aliquot of SCD submitted for PCR, it can be assumed that these discrepancies arose from the ability of specific nucleic acids to adhere to the CM, or not, as well as the sensitivity of the PCR assay for the specific agent. The most likely reason for the failure to detect these agents likely reflected poor adherence of the specific nucleic acid to the media. We presume CM’s enhanced effectiveness reflected the specific nucleic acid’s ability to adhere to the media, concentrating it above the detection limit of the assay. While individual assay failures could also account for false negative results, a control was used for each reaction.

Importantly, many organisms identified by PCR in SCD and/or the CM samples, including MuAstV1, MNV, MuCPV, SFB*, Chlamydia muridarum, S. xylosus, C. bovis, Helicobacter spp., Klebsiella oxytoca, Spironucleus muris*, and *Tritrichomonas* spp., are enzootic in most colonies in our vivaria. The study could not ascertain whether the source of nucleic acid for these agents were from NYC streets or vivaria floors as most users enter and exit the facilities multiple times daily using the shoe cleaner on each entry. Nucleic acid for 22 tested agents could only have originated from shoes contaminated with outdoor debris as these agents are excluded from our colonies. Most importantly, regardless of the nucleic acid source, none were infectious as none of the mice exposed to SCD and/or CM, or inoculated with extracts from these materials, became infected with any agent.

We routinely test wild mice captured in or near our vivaria to identify agents of concern to our research mouse populations. These mice frequently are positive for LDV, MCMV, MPV-1, MPV-2, MPV-5, MNV, MTV, MuAstV1, MuAstV2, MuCPV, murine picornavirus, and EDIM. Therefore, it was not surprising to find that the SCD and/or CM were positive for many of these agents. Interestingly, while MCMV, MTV, EDIM, LDV and murine picornavirus are commonly detected in wild mice, they were never detected in any SCD and/or CM sample. While endo and ectoparasites are among the most common contaminants of contemporary mouse colonies, it would appear unlikely that their introduction or spread is the result of contaminated footwear.^15,43,55^ While none of the SCD or Cm was shown to contain infectious material, the detection of ectromelia, MAV 1 & 2, MVM and MPV are of significant concern, as these agents are commonly excluded from research mouse colonies and both MAV 2 and MPV appear to be circulating within colonies of wild NYC mice.^56^

Some of the bacterial or protozoal agents assessed in this study for which nucleic acid was detected in the SCD and/or CM samples are excluded from all vivaria tested in this study. Therefore, as with some of the viruses, detection of these agents cannot be attributed with certainty to shoes contaminated with outdoor debris. Although we do not test and exclude sarbecovirus from our colonies, we assume it initially originated from outdoor debris, given that it was non-existent in the U.S. before 2020. SCD and/or CM samples from 3 vivaria tested positive for *Ornithonyssus bacoti*, an ectoparasite excluded from all vivaria. Surprisingly we did not detect other mites or nematodes in SCD or CM even though we frequently find murine pinworms (e.g., *Syphacia obvelata*) and fur mites (e.g., *Radfordia affinis*) in wild mice captured in or near our vivaria.

Additionally, although not specifically reported in NYC wild mouse or rat populations, rodents are known to carry other pathogens such as *Bordetella bronchiseptica, Campylobacter coli, Filobacterium rodentium, Francisella tularensis, Leptospira spp., Salmonella* spp., LCMV, *Eimeria* spp. and *Hymenolepis* spp.^1,18,20,23,24,39,48,53,55,56,57,59^ PCR results from SCD and CM were negative for all these agents suggesting a low biosecurity risk.

Interestingly, the control CM and SCD samples tested positive for several agents despite being autoclaved. Previous studies have shown that 15 minutes of autoclaving at 121°C and 15 psi is insufficient to degrade DNA and prevent nucleic acid re-amplification during PCR.^16^ Using these temperature and pressure parameters, it can take up to 2 hours of autoclaving to eliminate all nucleic acids.^21^ Our autoclave cycle consisted of a pulsed vacuum cycle with 4 pulses, a maximum pressure of 12 psi, and a temperature of 121°C (250°F) for 20 minutes. Therefore, it is not surprising that we detected nucleic acids in our samples. However, this is not consistent with previous work assessing infectious agent transmission from autoclaved cages.^10,42,47^

The most surprising and important aspect of this study was that SW and NSG mice directly inoculated with and housed in direct contact with SCD and/or CM exposed to SCD did not become colonized or infected with any of the agents found in SCD and/or CM with perhaps *Pseudomonas aeruginosa* being the sole exception. As mice utilized in this study were infected with MuAstV1, SFB and *S. xylosus* as these agents are enzootic in their colonies of origin, we could not assess whether SCD/CM could transmit these agents; however, based on the findings with numerous other agents detected in this study we believe their transmission would be highly unlikely.

Mice exposed to SCD from vivarium B were PCR positive for Beta *Streptococcus* Group B and *Hexamastix muris* in the pooled day 3 and 7 sample; however, the copy numbers were extremely low and the negative results from samples collected at later time points indicates these results were likely false positives. Similarly, at 63 DPI, mice in the Control group exposed to the autoclaved SCD inoculum tested positive for *Demodex* spp., *Demodex musculi*, and *Pseudomonas aeruginosa* by PCR. The copy numbers for *Demodex* spp., and *D. musculi* were low at 100 and 12, respectively. Although we have seen *D. musculi* transmission to soiled bedding sentinels in our health monitoring program, colonization typically occurs from dam to offspring, and mite numbers in immunocompromised mice are usually high enough to consistently identify positive results by skin scrapes and PCR.^38,50^ Considering the inoculum was autoclaved, the low copy number, and negative confirmatory tests on NSG mice (deep skin scrapes and skin biopsies), we consider the PCR result to most likely reflect a false positive result.

*P. aeruginosa* was also detected in the autoclaved Control SCD and vivarium D 63 I samples; however, the copy number for the control sample was very low (25), the mice had not tested positive at earlier time points, and the NSG mice were normal clinically at necropsy suggesting these results also likely reflect false positive results. Highly immunocompromised mice such as NSG, SCID and NOG are known to be highly susceptible to *P. aeruginosa*-induced disease.^14,41^ The copy number for vivarium D was 1622 suggesting the bacterium may have been present in the sample. However, none of the earlier samples were positive. This timing raises doubts as to whether infection resulted from exposure to SCD or, more likely, another source. *P. aeruginosa* is a common environmental opportunist found ubiquitously in soil and water, with horizontal transmission occurring within 5 days in mice.^30^

SW mice exhibited lymphoid and GALT hyperplasia at the final timepoint. GALT hyperplasia has been associated with *C. muridarum* infection.^37^ However, MOMP IHC staining of large intestinal tissue and testing of pooled lymph node biopsies by PCR were negative for the bacterium. We could not identify an infectious etiology. We speculate these lesions likely reflected antigenic stimulation following exposure to the SCD. All other microscopic findings were considered incidental, spontaneous, strain, or age-related, and were not associated with SCD exposure.^8,35,49^

As the vivarium shoe cleaner study results were unexpected, we inoculated additional NSG mice with pooled freshly collected and original sample SCD from all vivaria. These mice remained clinically normal and lesion free at 63 DPI. As the SCD used in the initial experiments had been collected over a period of up to 335 days, we also assessed footwear directly by collecting samples from the soles of animal care staff’s street shoes immediately on facility entry using CM and exposed mice directly to these CM samples. While nucleic acid from 11 agents was detected in an extract from the pooled CM sample, all NSG and SW mice exposed to contaminated CM samples were PCR negative at 63 DPI. However as discussed, the mice used in this study were MuAstV1 and SFB positive and therefore we could not assess whether these agents could be transmitted via contaminated footwear.

We encountered 6 PCR assays yielding inconclusive results preventing determination of the full range of infectious agents that may have been present in the samples. These agents of interest included: MPV4, SFB, *Campylobacter coli, Campylobacter jejuni, Helicobacter rodentium,* and *Streptobacillus moniliformis*.

Inconclusive results were obtained more frequently in SCD samples AS compared to CM, and when testing for bacteria as compared to viruses or parasites. However, if these agents were present and infectious we would have detected them in the samples collected from the exposed and inoculated mice.

In conclusion and importantly, these results demonstrate that while nucleic acid from many rodent pathogens can be found on footwear, it is unlikely footwear poses a biosecurity risk to research mouse colonies, questioning the necessity of not only the use of shoe cleaners, but also the use of shoe covers as a component of a rodent biosecurity program.

## Acknowledgements

The authors would like to thank Abigail Michelson for support during data collection, Simona Bekker and John D’Allara for their technical assistance during sample handling and shipment, and Fabricio Munoz and Marcia Lewis for their assistance with the breeding of the NSG mice used in this study. We also thank the Laboratory of Comparative Pathology staff for their technical assistance with histology and special stains.

## Conflict of Interest

The authors have no competing interest to declare.

## Funding

MSK Core Facilities are supported by MSK ’s NCI Cancer Center Support Grant P30 CA008748.

## Abbreviations

NYC: New York City
SCD: Shoe Cleaner Debris
CM: Contact Media
GEM: genetically engineered mouse
NSG: NOD.Cg-Prkdc^scid^ Il2rg^tm1Wjl^/SzJ
SW: Tac:SW
DPI: days post inoculation
OR: Odds Ratio
WBL: wire bar lid

## Notes

### Competing Interest Statement

The authors have declared no competing interest.

