## Supplemental table 1 for "Assessing the Biosecurity Risk of Footwear as a Fomite for Transmission of Adventitious Infectious Agents to Mice"

| **Supplemental Table 1.** Viral, Bacterial and Protozoa That Were Not Detected by Serology and/or PCR in Shoe Cleaner Debris (SCD) or Contact Media (CM) | | | | |
| --- | --- | --- | --- | --- |
|  | **Shoe Cleaner Study** |  | **Street Shoes Direct Testing Study** | |
| **Pathogen** | **SCD or CM^a^** |  | **CM exposed to Shoe Soles^b^** | **Mice exposed to CM exposed to Shoe Soles^b^** |
| Boone cardiovirus-1 | - |  | - | NP |
| **Hantaan virus** | - |  | - | - |
| New World Hantaviruses | - |  | - | - |
| Old World Hantaviruses | - |  | - | - |
| **LDV** | - |  | - | - |
| **LCMV** | - |  | - | - |
| **MCMV** | - |  | - | - |
| **MTV** | - |  | - | - |
| **EDIM** | - |  | - | - |
| Murine alphacoronavirus | - |  | - | NP |
| Murine kobavirus-1 | - |  | - | NP |
| Murine kobavirus-2 | - |  | - | NP |
| Murine picornavirus | - |  | - | NP |
| **MuCPV** | - |  | - | - |
| Murine sapovirus | - |  | - | NP |
| **PVM** | - |  | - | - |
| **Reovirus-3** | - |  | - | - |
| **Sendai virus** | - |  | - | - |
| **TMEV** | - |  | - | - |
| Group A *Streptococcus* | - |  | - | - |
| Group G *Streptococcus* | - |  | - | - |
| *Bordetella bronchiseptica* | - |  | - | - |
| *Bordetella pseudohinzii* | - |  | - | - |
| *Francisella tularensis* | - |  | - | NP |
| *Leptospira* spp. | - |  | - | NP |
| ***Mycoplasma pulmonis*** | - |  | - | - |
| *Rodentibacter pneumotropicus** | - |  | - | - |
| *Salmonella* spp. | - |  | - | - |
| *Cryptosporidium* spp. | - |  | - | - |
| ***Encephalitozoon cuniculi*** | NP |  | - | NP |
| *Giardia* spp. | - |  | - | - |
| Mite (*Myobia musculi*, *Myocoptes musculinis*, and *Radfordia affinis* | - |  | - | - |
| Pinworm (*Syphacia* spp. and *Aspiculuris* spp.) | - |  | - | - |
| *Pneumocystis* spp. | - |  | - | NP |
| Tapeworm | - |  | - | NP |

**Abbreviations:** NP, Not Performed

**Bolded agents were tested for by serology and negative.** Refer to Table 1 for full list of agents tested by serology

**^a^** Representative of negative results from Tables 3, 4, 5 or 6

**^b^** Representative of negative results from Table 7

* Enzootic in sampled vivaria. Other agents are either excluded or not monitored.
